# Erythropoiesis differences in various clinical phases of Dengue fever using Immature Reticulocyte Fraction parameter

**DOI:** 10.1101/563924

**Authors:** Amaylia Oehadian, Putri Vidyaniati, Jeffery Malachi Candra, Uun Sumardi, Evan Susandi, Bachti Alisjahbana

## Abstract

Dengue fever is the most common cause of hospitalization in otherwise healthy person. The incidence rate increases significantly every year, from 0.05 per 100.000 inhabitants in 1968 to 35-40 per 100.000 inhabitants in 2013. Thrombocytopenia and leucopenia are the most commonly reported changes, whether they are caused by bone marrow suppression or peripheral destruction. Anemia has also been observed to occur in dengue infection with unknown mechanism. Immature Reticulocyte Fraction (IRF) is a parameter reflecting the most immature reticulocyte fraction and it can identify the earliest stage of erythropoiesis disorder. This research aimed to determine the mechanism of erythropoiesis disorders that led to anemia using IRF parameter in the various clinical phases of dengue fever. This study was a comparative analytical research using secondary data derived from the *Dengue-associated Endothelial Cell Dysfunction and Thrombocyte Activation* (DECENT) research. The baseline characteristic data consists of sex, age, level of hemoglobin, hematocrit, leucocyte, and thrombocyte, also the IRF. The patients were grouped into the fever, critical, recovery, and convalescent phases, plus healthy control group. The data was analyzed using the Kolmogorov-Smirnov normality test, followed by Friedman test and Mann-Whitney post hoc test. There were 244 research subjects and the median age was 24 (14-67), with similar ratio of male and female. The median IRF for all the research subjects was 4.8% with an IQR of 2.4-8.1%. The fever-phase group showed a median of 1.8% with an IQR of 0.5-2.85%. The critical-phase group showed a median of 3.6% with an IQR of 1.8- 5.0%, while the median for the recovery-phase group was 7.05% with an IQR of 4.08-11.85%. The convalescent-phase group showed a median of 7.3 % with an IQR of 3.95-9.3%, and the healthy-control group showed a median of 4.1% with an IQR of 2.2-6.6%. There was a significant difference in IRF between the groups (p<0.05). Immature Reticulocyte Fraction in fever phase was significantly different with IRF in other phases and healthy controls (p<0.05). In the critical phase, IRF was not significantly different from the healthy controls (p=0.218). Summarizing, there are changes in erythropoiesis activity in various clinical phases of dengue infection. Erythropoiesis suppression occurred mainly during the fever phase and it started to restore in the critical phase. In the recovery and convalescent phases, the erythropoiesis activity was increased.

## Introduction

Dengue infection is the most notable viral infection transmitted by mosquitoes, seen from the medical perspective and also the community health perspective. The incidence of dengue infection has increased significantly every year, from 0,05/100.000 in 1968 to 35-40/100.000 in 2013 [1,2].

The hematological disorders known in dengue viral infection are temporary thrombocytopenia and leucopenia. The mechanism of neutropenia during dengue infection could be caused by bone marrow suppression or peripheral destruction. Erythropoiesis disorder is also observed in dengue infection, which maybe caused by the bone marrow suppression affecting all of the hematopoiesis series. It is hypothesized that a combination of viral infection on hematopoietic progenitor cells, and the viral infection on bone marrow stromal cells and dengue specific T-cell activation, both releasing cytokines that suppress the bone marrow [3,4].

Erythropoiesis suppression during dengue infection was also shown in the form of aplastic anemia by several case reports. Khoj et al reported aplastic anemia occurred after dengue infection and ended with bone marrow transplantation [5]. A study by Lora et al in Dominican Republic indicated that anemia related to the severity and mortality of dengue infection, found in 19% of patients and 32% of severe dengue infection with an Odd Ratio of 3 for mortality [6]. Clinical manifestation of dengue infection in thalassemic patients was different, where they experienced the decrease of hemoglobin rather than hemoconcentration [7].

Erythropoiesis disorder in dengue infection has not been extensively studied. Bone marrow examination is an invasive procedure and not recommended in dengue infection. A simple hematological parameter such as *Immature reticulocyte fraction* is expected to be able to indicate how active the erythropoiesis in the bone marrow is [8].

*Immature reticulocyte fraction* (IRF) is a parameter reflecting the most immature reticulocyte fraction. This IRF parameter is simple and can be obtained directly from the automated hematology analyzer Sysmex XE-2100. The IRF reflects erythropoiesis directly and identifies erythropoiesis disorder earlier than reticulocyte and hemoglobin. A study from Goncalo in hematological malignancy patients undergoing hematopoietic progenitor cells transplantation showed IRF as an earlier indicator for success than neutrophil and thrombocyte with two to four days of difference [8–10]. We conduct this study in order to observe whether erythropoiesis is really suppressed during Dengue infection.

## Method

This is a cohort study, using data derived from the *Dengue-associated Endothelial Cell Dysfunction and Thrombocyte Activation* (DECENT) research in the Department of Internal Medicine which ran from March 2011-March 2012. The DECENT study was designed as a cohort study that recruited patients presenting with clinical signs of symptomatic Dengue virus infection (SDVI) to Hasan Sadikin Hospital in Bandung. Consecutive patients meeting inclusion criteria were enrolled and followed with daily clinical assessment, blood collection and, in a subgroup; assessment of plasma leakage until day 5 post-admission. Temporal changes on laboratory (thrombocytes and endothelial cells) parameters and plasma leakage during the infection were determined. Patients were asked to return for follow-up blood collection at >2 weeks (14-20 days) after discharge. Inclusion criteria are subject must be 14 years old or above, and clinical suspicion or confirmation of having DF or DHF/DSS according to WHO criteria. Exclusion criteria are pregnancy, clinical symptoms/signs of or known malignancy, known coagulation disorder, and any chronic diseases such as diabetes mellitus, chronic renal failure, hepatitis, auto-immune disorders and psychiatric disorders.

The statistical test using the Friedmann test was used to examine the difference between the phases in dengue patients (4 groups), followed by a *post hoc* Mann Whitney test, the result is considered to be significantly different if the p value is <0.05 [11].

## Results and discussion

### Results

#### Baseline characteristics of research subjects

There were 244 research subjects with a similar ratio of male and female, 56.6% of males and 43.4% of females, with the median age of 24 and IQR (interquartile range) of 14-67. The hemoglobin and hematocrit levels were normally distributed in every phase with a mean of 14.0 ± 1.9 g/dL for hemoglobin and 41.1 ± 5.2 % for hematocrit. Whereas, the leucocyte and thrombocyte levels were not normally distributed, with the leucocyte median of 5.100/mm^3^ and IQR of 3.800/mm^3^ to 6.600/mm^3^, and the thrombocyte median of 78.000 mm^3^ and IQR of 41.000/mm^3^ to 177.000/mm^3^. The baseline characteristics for each phase and control group can be seen below (Table 1).

**Table 1.** Baseline Characteristics of Research Subjects.

| Variable | Clinical phase of dengue infection |  |  |  | Healthy control<br>n=27 | p* value |
| --- | --- | --- | --- | --- | --- | --- |
|  | Fever phase<br>n=13 | Critical phase<br>n=77 | Recovery phase<br>n=74 | Convalescent phase<br>n=53 |  |  |
| Sex (n, %) |  |  |  |  |  |  |
| Male | 8 (61.5) | 46 (59.7) | 42 (56.8) | 32 (60.4) | 10 (37.0) | 0.292<sup>c</sup> |
| Female | 5 (38.5) | 31 (40.3) | 32 (43.2) | 21 (39.6) | 17 (63.0) |  |
| Age (year)<br>median<br>IQR | 24<br>(20-28) | 24<br>(19-33) | 23<br>(19-33) | 25<br>(20-34) | 25<br>(22-31) | 0.810<sup>b</sup> |
| Hb (g/dL)<br>mean<br>± SD | 13.8<br>± 2.4 | 14.5<br>± 2.1 | 13.7<br>± 1.7 | 13.9<br>± 1.7 | 14.2<br>± 1.8 | 0.076<sup>a</sup> |
| Ht (%)<br>mean<br>± SD | 40.0<br>± 6.1 | 42.2<br>± 5.8 | 39.9<br>± 4.8 | 41.5<br>± 4.3 | 42.3<br>± 4.7 | 0.040<sup>a</sup> |
| Leucocyte<br>(x1000/mm<sup>3</sup>)<br>median<br>IQR | 3.3<br>(2.4-4.3) | 4.0<br>(3.0-5.6) | 5.0<br>(4.0–6.0) | 7.1<br>(5.7-8.8) | 7.0<br>(6.5-8.0) | <0.001<sup>b</sup> |
| Thrombocyte<br>(x1000/mm<sup>3</sup>)<br>median<br>IQR | 66<br>(43-89) | 36<br>(19-55) | 94<br>(57-132) | 265<br>(225-313) | 293<br>(250-318) | <0.001<sup>b</sup> |

\*Note : analysis using a: ANOVA test, b: Friedman test, c: Chi Square test, significant if p<0.05

#### The IRF differences among dengue infection phases

The IRF differences in various clinical phases of dengue infection and control group can be seen in Table 2 below. The median IRF for all the research subjects was 4.8% with an IQR of 2.4-8.1%. The fever-phase group showed a median of 1.8% with an IQR of 0.5-2.85%. The critical-phase group showed a median of 3.6% with an IQR of 1.8-5.0%, while the median for the recovery-phase group was 7.05% with an IQR of 4.08-311.85%. The convalescent-phase group showed a median of 7.3 % with an IQR of 3.95-9.3%, and the healthy-control group showed a median of 4.1% with an IQR of 2.2-6.6%.

**Table 2.**
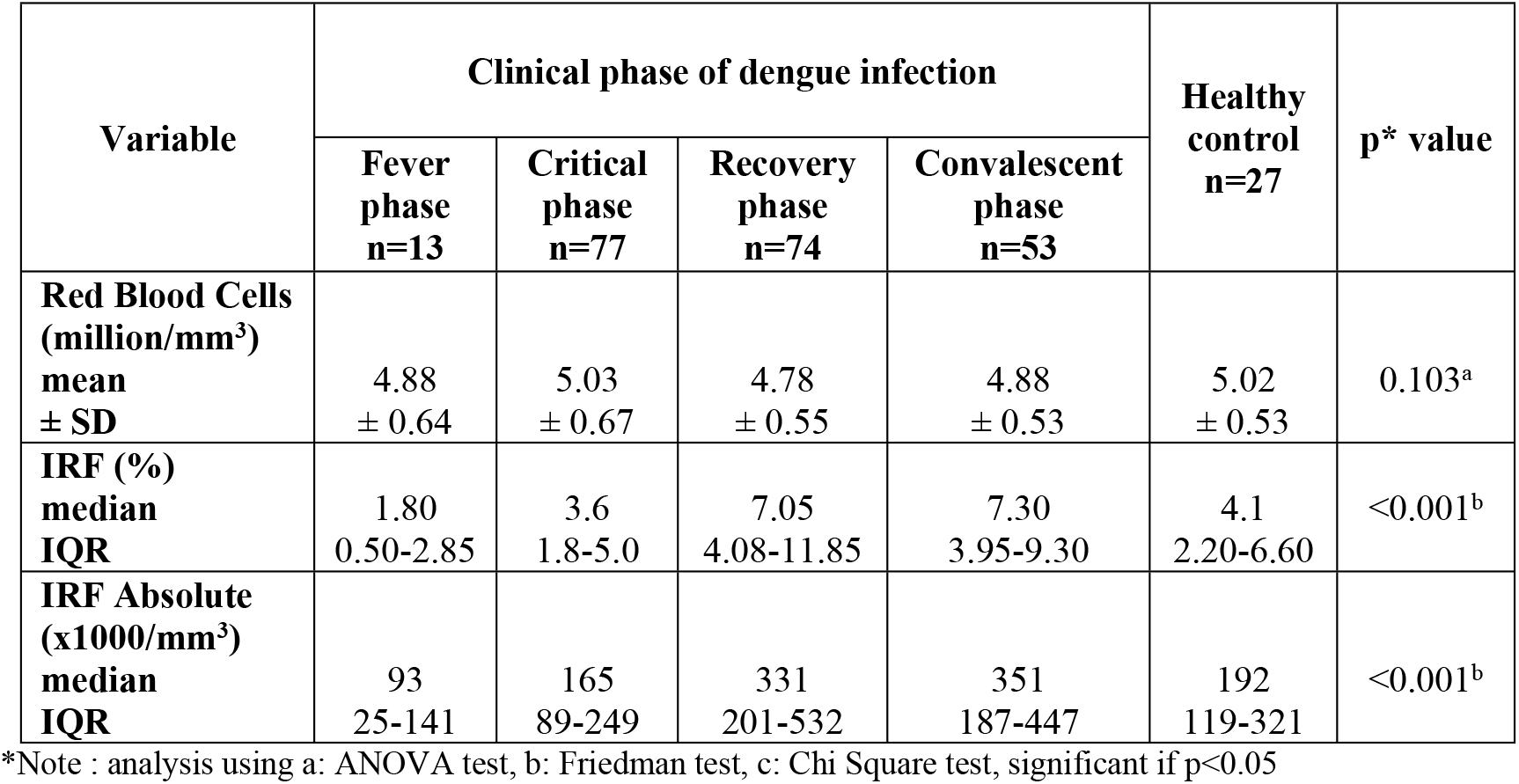
Red Blood Cell Quantity and Immature Red Cell Fraction in Various Phases of Dengue Infection.

The result of the statistical test using the Friedman test on IRF (%) in various clinical phases of dengue infection showed a significant difference with a p value of < 0.001. The statistical analysis was then followed by a *post hoc* Mann Whitney test to examine the difference between each group, and the difference is considered to be significant if p<0.05 (Fig 1).

**Fig 1.**
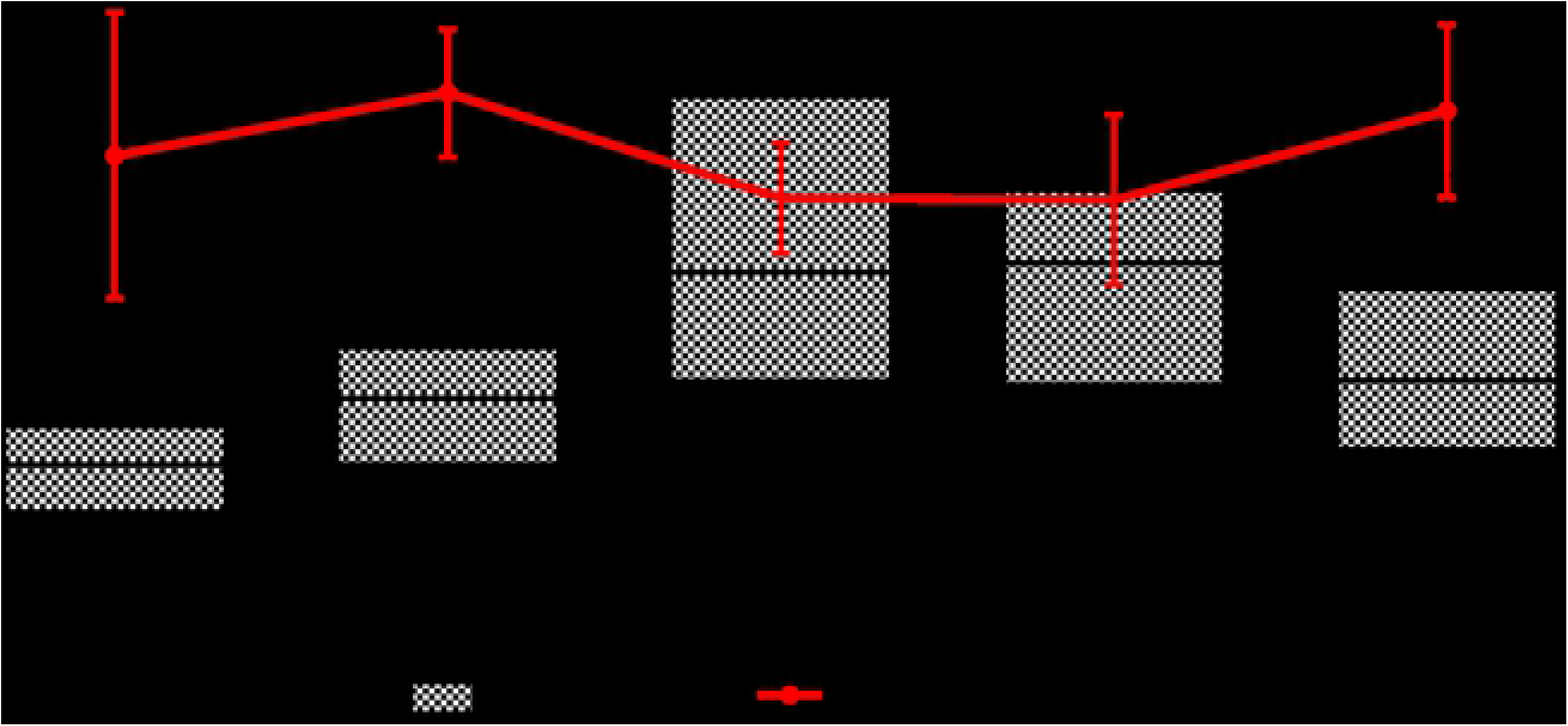
Scattergram of IRF in Clinical Phases of Dengue Infection (please find attached “IRF rev”, containing the figure)

The IRF (%) was lowest in the fever phase, then increased in the critical phase, reaching its highest in the recovery phase, and decreased in the convalescent phase. The IRF (%) in the fever phase was significantly different when compared to all the other phases and the healthy control group, with all the p value < 0.05. The IRF (%) in the critical phase was significantly different when compared to the other phases in dengue infection (p < 0.05), but not when compared to the healthy control group (p = 0.218). The IRF (%) in the recovery phase was significantly different when compared to the fever phase, critical phase, and the healthy control group, but not when compared to the convalescent phase (p = 0.629). The IRF (%) in the convalescent phase was significantly different when compared to the fever phase, critical phase, and the healthy control group (p < 0.05). When compared to IRF (%), RBC counts were not significantly different among the phases of clinical dengue and also healthy control group.

## Discussion

### Baseline characteristics of research subjects

There were a total of 244 research subjects, with the median age of 24 and IQR (interquartile range) of 14-67. The number of male and female subjects was similar. The age and sex of the subjects in this reasearch were consistent with the report from *Pusat Data dan Surveilans Kementrian Kesehatan Republik Indonesia* in 2010 stating that cases of dengue infection in Indonesia occurred predominantly in the age group of more than 15 years old with the distribution of the cases nearly the same in both sexes [1].

The hemoglobin levels in this research were not significantly different among the groups. A study by LaRussa concluded that the bone marrow suppression in dengue infection happened so fast, that even though the bone marrow suppression affected all hematopoietic series, the substantial erythrocyte reserve can compensate for the decrease in erythrocyte production [3].

The result of the statistical test on hematocrit levels in every group showed a significant difference between the groups, with the highest hematocrit level in the critical-phase group. The increase in hematocrit levels corresponds to the hemoconcentration and plasma leakage that happens during the critical phase. Rothman and Malasit suggested that the increase in capillary permeability was caused by the involvement of mediators such as TNF-α, IL-2, IL-8 and VEGF that would disrupt endothelial cells permeability in vitro. The intensity of the immune response and the titer of plasma viremia are the strongest independent factors for plasma leakage [12–13].

The leucocyte counts among the groups were significantly different. The results showed that the lowest leucocyte count was found in the fever-phase group, then increased gradually according to the disease phase. A study by Simmons, Na-Nakorn and LaRussa stated that leucopenia and neutropenia were found in the early phase of the disease, and a bone marrow biopsy in the early phase of the disease (less than five days of fever) also showed more hypocellularity [3,14].

The statistical test on the variable of thrombocyte level showed significant difference between the groups. The thrombocyte level started decreasing in the fever phase, with the lowest level found in the critical phase and started increasing again in the recovery and convalescent phases. The result was consistent with the pathogenesis of dengue viral infection where the suppression of bone marrow happens in the early fever phase and reaches its lowest point in the critical phase [3].

### The IRF differences among dengue infection phases

The IRF in the fever phase was the lowest compared with other phases of dengue infection. The IRF started rising in the critical phase and kept rising in the recovery and convalescent phases.

The IRF in the fever-phase group was significantly different compared with the other groups. The low IRF reflected the bone marrow suppression occuring in the fever phase of dengue infection. This result was consistent with the bone marrow biopsy research done by Simmons et al showing that the onset of bone marrow suppression could happen not more than 12 hours after the infection. A study by Noisakran also showed that the dengue virus can reach the bone marrow compartment in a short period of time, confirming that bone marrow suppression occurs in the fever phase [3,15].

The analytical result of IRF in the critical phase showed a significant difference compared with other groups, except the healthy-control group. This showed that the recovery of bone marrow suppression has started in the critical phase. A study by Noisakran showed that the number of dengue virus RNA reached a peak in day 1 to day 3 after the infection and then the number decreased, so the bone marrow suppression by dengue virus RNA lessened in the critical phase (day 4 of sickness). LaRussa and Noisakran concluded that bone marrow recovery ended in day 10 (afebrile period) even though the blood cell destruction (especially thrombocyte) in the periphery reached its peak (owing to antibody and complement clearance) [3,15]. A research by Na-Nakorn also stated that a bone marrow examination in day 4 to day 8 (critical phase) of sickness showed erythroid hyperplasia with maturation disruption [14].

The IRF in the recovery-phase group was not significantly different compared to the convalescent-phase group. The IRF in the recovery-phase group and convalescent-phase group were higher than the critical-phase group and the healthy-control group. This result was similar with a bone marrow examination by Na-Nakorn in day 10-14 of sickness that showed erythropoiesis hyperplasia without maturation disruption [14].

The analytical result of IRF in the critical-phase group and healthy-control group using the Mann-Whitney test did not show any significant difference. The pathogenesis of dengue infection states that thrombocyte and leucocyte reach the lowest level in the critical phase, but the same thing did not happen to the erythrocyte in this reasearch, represented by the IRF. This result was similar to research by LaRussa and Na-Nakorn which showed that bone marrow suppression started its recovery in the critical phase, characterized by erythropoiesis hyperplasia with maturation disruption [3,14-15]. The increase in erythropoiesis is usually followed by an increase in erythropoietin. The life span of reticulocytes in the circulation also increases for up to three days or even more, due to the release of ‘stress or shift’ reticulocytes from the bone marrow and the acceleration of erythroid differentiation. ‘Stress or shift’ reticulocytes from the bone marrow have a large amount of RNA in their cells. The population of existing reticulocytes can not only be counted, the RNA content according to its maturity level needs to be assessed as well, and this has a significant clinical application in evaluating the erythropoiesis activity [16].

The comparison between IRF in the critical-phase group and healthy-control group did not show significant difference, which shows that bone marrow of dengue patients in the critical phase is not different from the bone marrow of healthy people in general. This result confirmed that thrombocytopenia occuring in critical phase of dengue infection is caused by cell destruction in the periphery, due to the antibody and complement clearance [3].

## Limitation of Research

This research has a few limitations, which are:

1. The comparison in the number of samples between the fever-phase group and the critical-phase group with the other groups are very different,
2. The method of this research was prospective, so the data is dependent on the existing records.

## Conclusion

This research shows that there is an erythropoiesis difference between the phases of dengue infection. Bone marrow suppression for erythropoiesis reaches the lowest point in the fever phase and starts recovering in the critical phase. Erythropoiesis in the critical phase is not different from the healthy control. This study was also the first study describing IRF in multiple phases of dengue disease.

